# Local states of chromatin compaction at transcription start sites control transcription levels

**DOI:** 10.1101/782060

**Authors:** Satoru Ishihara, Yohei Sasagawa, Takeru Kameda, Mana Umeda, Hayato Yamashita, Naoe Kotomura, Masayuki Abe, Yohei Shimono, Itoshi Nikaido

## Abstract

A defined amount of transcript is produced from transcription start sites (TSSs) of each gene, suggesting that the binding frequency of RNA polymerase varies among genes. However, what structure in chromatin controls this frequency remains elusive. We established a method to fractionate chromatin according to its degree of three-dimensional compaction. Nucleosomes were evenly detected through all of the fractions, but histone H1 was more highly enriched in the more compact chromatin fractions. Similarly, HP1α and MBD2b were more abundant in more compact chromatin, while the levels of tri-methylated H3 (Lys9) and 5-methyl cytosine subtly increased. Via genome-wide analyses, nearly the entire genome was found to exist in compact chromatin without variations between repeat and non-repeat sequences; however, active TSSs were rarely found in compact chromatin. Based on a correlation between weak compaction and RNA polymerase binding at TSSs, it appears that local states of chromatin compaction determine transcription levels.

## INTRODUCTION

All physiological reactions on the genomic DNA require the binding of protein factors with various enzymatic activities to the appropriate regions of the genome. However, these reactions occasionally fail when the protein factors cannot access their target regions. In eukaryotic cells in early S-phase, replication is initiated from replication origins within genomic regions where DNA-processing enzymes, such as endonucleases, are able to access the DNA under experimental conditions (*1, 2*). Repair and recombination in such enzyme-accessible regions also dominantly occur compared with the levels of these activities in enzyme-inaccessible regions (*3–5*). Similarly, more abundant transcripts are produced from genes within enzyme-accessible regions (*6–8*). These observations indicate that the accessibility of the genome directly controls the strength of reactions on the genome. Importantly, while replication, repair, and recombination are completed in a single round because of their all-or-nothing output, transcription is a repeated reaction because it is required to synthesize multiple RNA copies. The number of copies is determined by the frequency of the reaction, i.e., the frequency of RNA polymerase binding to the transcription start site (TSS). Therefore, a structure that can vary in its degree of accessibility is required to allow variations in transcription levels.

The chromatin structure largely influences the accessibility of the genome. As the primary structure of chromatin, the genomic DNA is wrapped around a histone octamer to form a nucleosome (*9*). The nucleosomes interact with neighboring nucleosomes and/or non-histone proteins to form a higher-order structure (*10*). Thus, the structures organized in this step-by-step process can hide the genomic DNA from protein factors and reduce the accessibility of the genome. Via treatment with micrococcal nuclease (MNase), the nucleosome positioning along the genome has been characterized. This investigation revealed that nucleosomes are absent in the region just upstream of the TSSs of actively transcribed genes (*11*). It is widely accepted that such regions, known as nucleosome-free regions (NFRs), support RNA polymerase binding (*11*), resulting in a direct correlation between chromatin structure and transcriptional state. However, because this level in the hierarchy of chromatin structure exists in two states, i.e., with or without nucleosomes, the ability to achieve intermediate levels of transcription might not be shown. Although NFRs are more clearly observed at the TSSs of highly transcribed genes via well-positioning of nucleosomes on both sides of the NFR (*12–15*), it has also been reported that the nucleosome positioning around the TSSs of genes that are uniformly transcribed in a given cell population is heterogenous (*16, 17*). In addition to nucleosome positioning, other variables are required in the hierarchy of chromatin structure to allow tuning of the transcription levels.

A nucleosome can interact with another nucleosome via the basic tail of histone H4, which has an affinity to an acidic patch on the surface of an adjacent nucleosome (*18, 19*). Therefore, an array of multiple nucleosomes, known as “beads on a string”, is often formed via inter-nucleosomal interactions to generate a more compact structure. A 30 nm thick fiber has been historically proposed to represent this more compact structure (*20*). Electron microscopy analyses have revealed that *in vitro*-reconstituted nucleosome arrays fold into a 30 nm thick rod-shaped structure that has been conceptualized by two alternative models: a one-start solenoid (*21*) or a two-start zigzag (*22*). Over the last decade, the results of imaging studies have argued against the existence of the 30-nm fiber and have instead revealed granular structures that are distinct from the rod of the 30-nm fiber (*23–25*). Recently, granules of various sizes were observed in nuclei of mouse embryonic stem cells, and these granules were estimated to consist of 4−8 nucleosomes (*26*). A derivative of the chromosome conformation capture (3C) method also revealed that motifs consisting of 3−4 nucleosomes were commonly formed in budding yeast (*27*). These observations together with live imaging analyses that showed fluctuating movement in individual nucleosomes (*26, 28–30*) suggest that a static 30-nm fiber structure is unlikely to be formed *in vivo*. How nucleosome arrays are in fact arranged remains under debate; nevertheless, it is not doubted that neighboring nucleosomes are locally compacted. Thus, such a structure must be responsible for regulating the accessibility of the genome, which ultimately controls transcription levels.

In this study, the three-dimensional compaction state within several neighboring nucleosomes was focused on as a possible chromatin structure responsible for controlling transcription levels. Via sedimentation velocity centrifugation, chromatin from cultured cells was successfully fractionated based on its local compaction state: compact chromatin sedimented into the lower fractions, while open chromatin remained in the upper fractions. Analyses of the fractionated proteins indicated enrichment of HP1α, MBD2b, and histone H1 in the lower fractions, suggesting that these proteins contribute to local chromatin compaction. Next-generation sequencing (NGS) of the DNA recovered from each fraction showed that nearly the entire genome was packaged into compact chromatin, with the exception of the chromatin at active TSSs, which was poorly compacted. The weakness of compaction at the TSS chromatin was more clearly correlated with transcription levels than was NFR formation. Together with a correlation with the RNA polymerase binding levels, the local state of chromatin compaction appears to influence the frequency of RNA polymerase binding, which ultimately regulates the transcription levels of individual genes.

## RESULTS

### Chromatin was fractionated according to its local compaction state

Chromatin is compacted via direct interactions among neighboring nucleosomes and other chromatin-related proteins. Once chromatin is treated with formaldehyde, such interactions are sufficiently preserved to examine the local state of chromatin compaction. For this study, we chose to use human hepatoma HepG2 cells, for which data on epigenetic marks are available in the ENCODE database (https://www.encodeproject.org/). HepG2 cells were mildly treated with 0.5% formaldehyde. Following solubilization with a detergent, the cells were sonicated to fragment chromatin, resulting in shearing of the genomic DNA into fragments of 300−500 bp on average. Next, the cell extract containing the fragmented chromatin was subjected to ultracentrifugation in a sucrose density gradient to fractionate the chromatin based on its degree of compaction (*31, 32*). Following removal of the uppermost layer (Fr-0), five fractions were collected and numbered Fr-1 to Fr-5, from the top to the bottom of the centrifugation tube, respectively (Fig. 1A). DNA was purified from each fraction and analyzed by agarose gel electrophoresis (Fig. 1B). The DNA appeared as smeared bands in all of the fractions, indicating that the chromatin was distributed throughout the centrifugation tube. Note that the formaldehyde concentration directly influenced the fractional distribution of the chromatin, i.e., when the formaldehyde concentration was reduced, the chromatin tended to remain in the upper fractions, possibly because of weaker preservation of the chromatin compaction (fig. S1). Because the average size of the DNA fragments in Fr-1 to Fr-5 was approximately 300−500 bp (Fig. 1B), arrays consisting of a few nucleosomes were mostly distributed in Fr-1 to Fr-5. Next, the proteins in each fraction were recovered by trichloroacetic acid (TCA) precipitation and analyzed by sodium dodecyl sulfate-polyacrylamide gel electrophoresis (SDS-PAGE). After reversal of the crosslinking via incubation at 65°C for 24 hrs, the proteins were loaded into an SDS-PAGE gel (Fig. 1C). Fr-1 to Fr-5 contained relatively fewer proteins, although an abundant amount of DNA was detected in Fr-1 to Fr-4 (Figs. 1C vs. 1B). To analyze the distributions of core histone H3 and linker histone H1, western blotting experiments with pan antibodies against each histone were performed. The protein preparations were normalized among the fractions based on the amount of DNA. There was slightly less H3 in Fr-1 than in the other fractions, and Fr-2 to Fr-5 contained similar levels of H3, indicating that a comparable number of nucleosomes were incorporated per unit length of DNA (“H3” in Fig. 1D and solid line in Fig. 1E). H1 was gradually enriched toward Fr-5, and it was estimated that the H1 abundance in Fr-5 was 3.2-fold (1.7 in log2) higher than that in Fr-1, indicating that H1 is involved in local chromatin compaction (“H1” in Fig. 1D and dotted line in Fig. 1E).

**Fig. 1.**
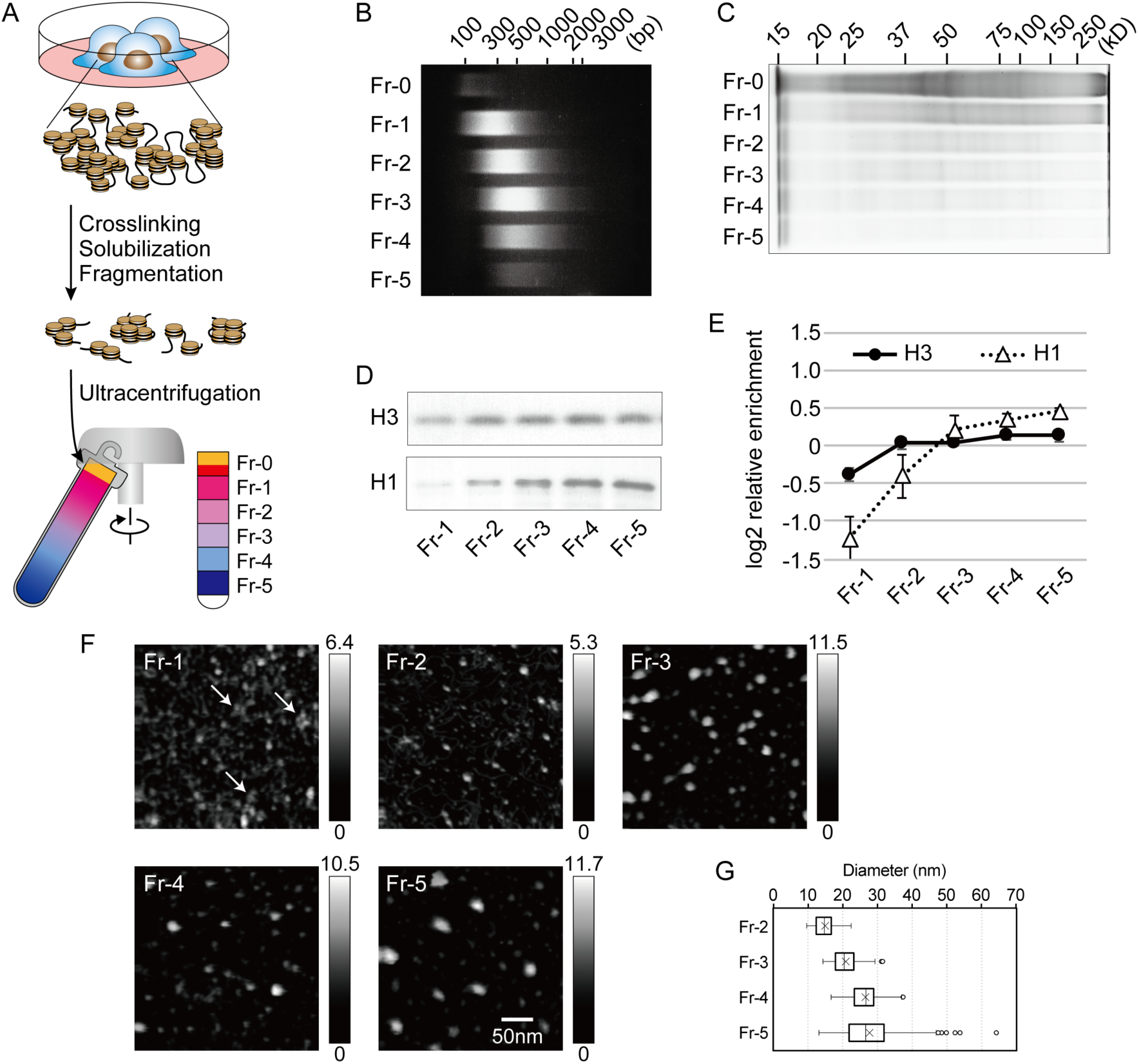
Chromatin fractionation by using sedimentation velocity centrifugation. (**A**) A schema of the fractionation method established in this study. (**B**) The size distribution of the DNA fragments in the fractionated chromatin. The DNA was size-separated on a 2% agarose gel. (**C**) The size distribution of the proteins in the fractionated chromatin. The proteins were size-separated on an 8% SDS-PAGE gel. (**D**) Western blotting for histones H3 and H1. The loaded samples were normalized among the fractions based on the amount of DNA. (**E**) The fractional distributions of histones H3 and H1. The relative enrichment of the histones in each fraction was calculated from the intensity of the blot signals and is represented by the log2 ratio to the average. Data obtained from at least three independent experiments are represented as the mean ± SD. (**F**) HS-AFM images of the fractionated chromatin. The arrows in Fr-1 highlight several closely gathered dots. The heights of the objects are shown as a grayscale gradation ranging from 0 to a maximum (nm) in each panel. All panels are shown at the same magnification. (**G**) The diameters of the chromatin fragments in each fraction. Data were obtained as described in the MATERIALS AND METHODS.

Chromatin was prepared from each fraction by immunoprecipitation with a pan anti-histone H3 antibody and then observed via high-speed atomic force microscopy (HS-AFM) (Fig. 1F). In Fr-1, objects including histone H3 were unstructured, although some dots did appear to be gathered (arrows in “Fr-1”). On the other hand, bright particles were observed in Fr-2 to Fr-5. Measurement of the particle diameters revealed that the particles became larger toward Fr-5, with the median diameters ranging from 15 nm (Fr-2) to 27 nm (Fr-5) (Fig. 1G). Because the length of the fractionated DNA was comparable among the fractions (Fig. 1B), the larger particles likely consisted of multiple arrays of nucleosomes and other chromatin-associated proteins. Thus, by using our fractionation method, chromatin was successfully fractionated according to its degree of local compaction.

### The distribution of epigenetic marks and readers in fractionated chromatin

Epigenetic marks are well known to influence chromatin structure. To evaluate the contribution of the marks to the local state of chromatin compaction, we performed immunoblotting analyses to investigate the fractional distribution of histone H3 modified at lysine residues in its amino-terminal tail. The loaded samples were normalized among the fractions based on the amount of DNA. When histone H3 lysine 9 was examined, unmodified and acetylated forms were evenly distributed throughout the fractions (“H3K9un” and “H3K9ac”, respectively, in Fig. 2A). On the other hand, tri-methylated H3 at lysine 9 was slightly enriched in Fr-2 to Fr-5 compared with its level in Fr-1 (“H3K9me3” in Fig. 2A and solid line in Fig. 2C). Similarly, a slight enrichment of tri-methylated H3 at lysine 27 was observed in the lower fractions (“H3K27me3”), but the levels of the unmodified and acetylated forms did not change (“H3K27un” and “H3K27ac”, respectively, in Fig. 2A). These observations suggest that methylation of these lysine residues is related to the local chromatin compaction, while acetylation appeared to be independent of the compaction. In addition, when cytosine methylation in CpG dinucleotides was assessed by dot blotting with an anti-5-methyl cytosine antibody, the signals were slightly increased toward Fr-5 (“5meC” in Fig. 2A and solid line in Fig. 2D). We further investigated the fractional distribution of “readers” of these epigenetic marks. The abundance of HP1α, a reader of H3K9me3, was increased toward Fr-5; however, the abundance of Suz12, a reader of H3K27me3, was decreased toward Fr-5 (“HP1α” and “Suz12” in Fig. 2B). These observations indicate that HP1α, but not Suz12, is preferentially present in compact chromatin. When the levels of MBD2 and MeCP2 (5meC readers) were examined, only the abundance of a small variant of MBD2, designated as MBD2b (*33*), was increased toward Fr-5 (“MBD2” and “MeCP2” in Fig. 2B). When the distributions of HP1α and MBD2b were compared with those of H3K9me3 and 5meC, respectively, both readers were more highly enriched toward Fr-5 relative to the levels of the epigenetic marks (dotted vs. solid lines in Figs. 2C and 2D). These patterns suggest that the readers are only recruited to a fraction of the available epigenetic marks before incorporation into compact chromatin.

**Fig. 2.**
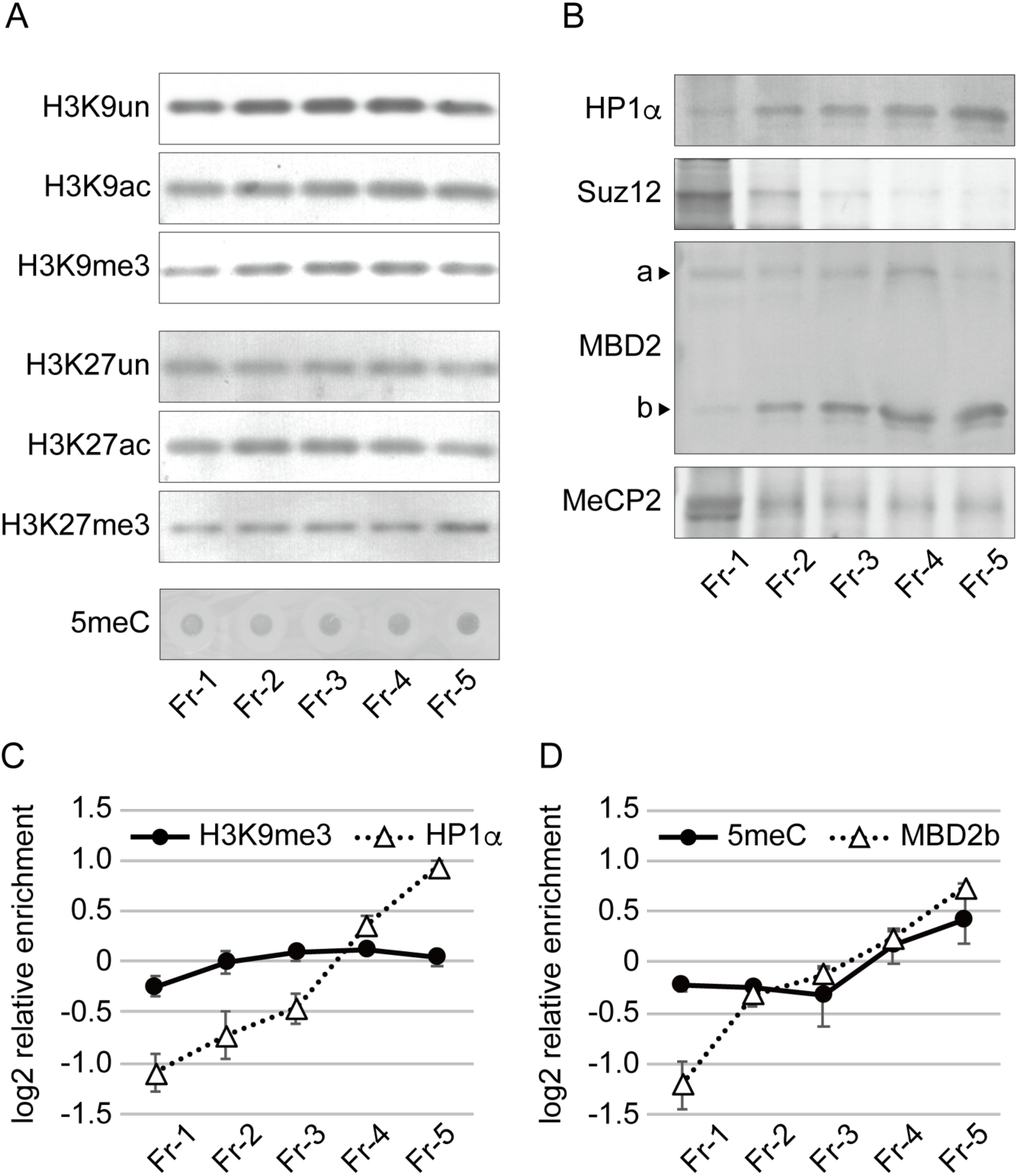
Immunoblot analyses of epigenetic marks and readers in fractionated chromatin. (**A**) The fractional distribution of the epigenetic marks was analyzed by western blotting. For 5meC, DNA from the fractionated chromatin was spotted onto a membrane and analyzed by dot blotting. (**B**) The fractional distribution of the epigenetic readers was analyzed by western blotting. In the “MBD2” panel, the large and small variants of MBD2 are marked by arrowheads labeled “a” and “b”, respectively. (**C** and **D**) The fractional distributions of the epigenetic marks vs. those of the readers. The relative enrichment of H3K9me3 and HP1α (C), and 5meC and MBD2b (D) was calculated from the intensity of the blot signals and is represented by the log2 ratio to the average. Data obtained from at least three independent experiments are represented as the mean ± SD.

### The distributions of active TSSs and repeat sequences in fractionated chromatin

Chromatin at TSSs of highly transcribed genes is opened via nucleosome eviction (*11*), while repeat sequences, such as transposon-derived elements, are mostly packaged into heterochromatin, which is well known to be a compact structure (*34, 35*). To investigate how these genomic regions were fractionated by our method, their distributions were assessed using quantitative PCR (qPCR) of DNA recovered from the fractions (Fig. 3A). More than 50% of the TSSs for the *GAPDH* and *ACTB* genes, which are abundantly expressed in HepG2 cells, was found in Fr-1, and the proportion gradually deceased toward Fr-5, although 2% of the signal was still detected in Fr-5. As representative repeat sequences, the distributions of *Alu*, L1, and α-satellite sequences were examined by qPCR with primer pairs that annealed to conserved regions in each repeat. The proportions of these repeat sequences in Fr-1 to Fr-3 ranged from 20% to 30%, although they were lower in Fr-4 (12−15%) and Fr-5 (8−9%). These values were quite similar to the proportions of total genomic DNA in the fractions. This similarity is attributable to the fact that almost half of the human genome consists of such repeats. Note that when a different formaldehyde concentration was used, the fractional distributions of the specific genomic regions changed (fig. S2). This outcome mirrors the effect on the distributions of the total genomic DNA (fig. S1). To simply describe the local state of chromatin compaction, log2 ratios of the proportions in Fr-5 and Fr-1 were calculated (“Fr-5/Fr-1” in Fig. 3B). The TSSs of the *GAPDH* and *ACTB* genes showed values of −5.36 and −4.65, respectively, while the values for the repeats and total DNA ranged from −1.79 to −1.25. The apparent differences between the active and repressed reference sequences indicate that the Fr-5/Fr-1 scores are useful for representing the relative levels of the local chromatin compaction.

**Fig. 3.**
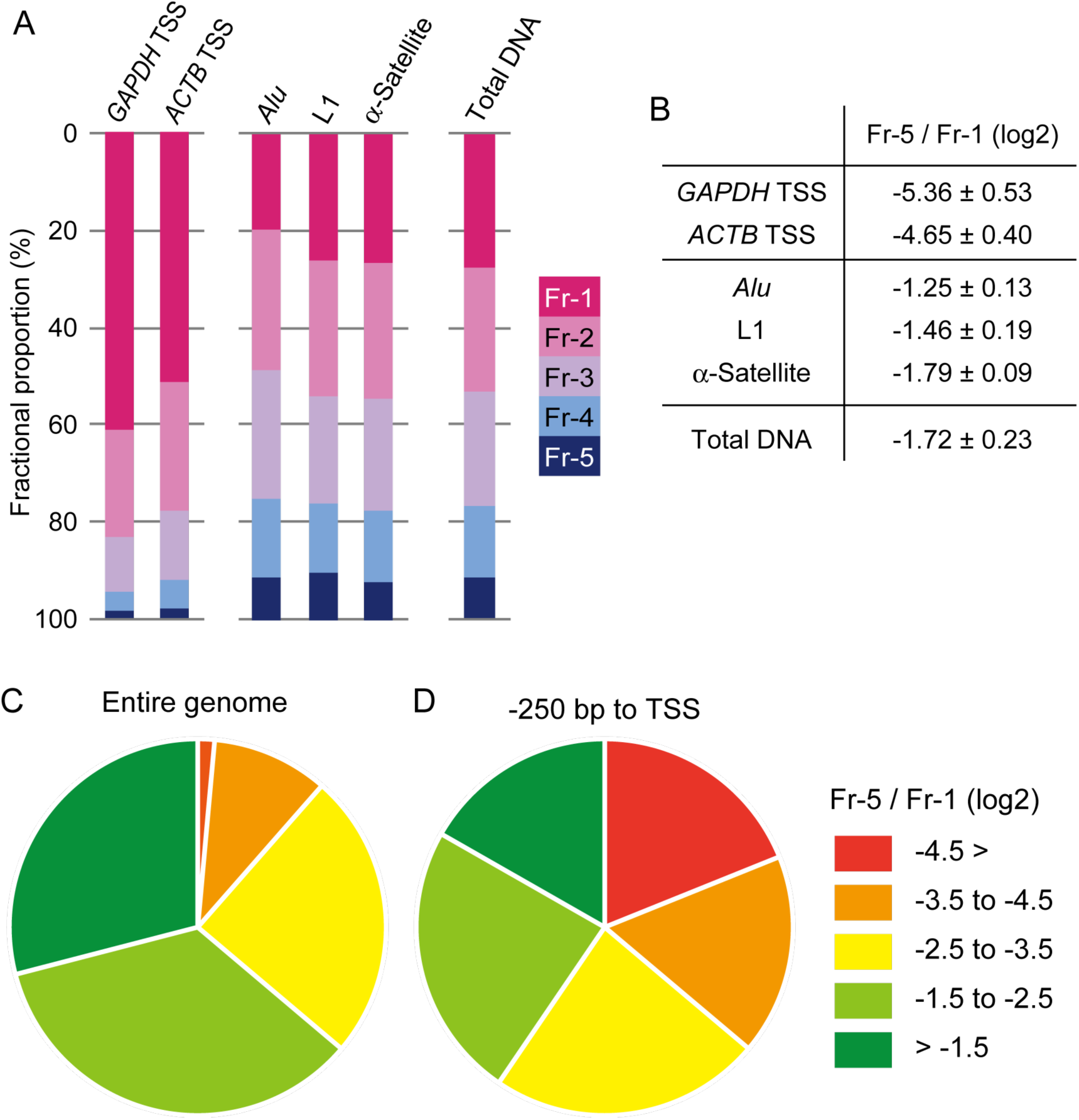
The fractional proportions and the Fr-5/Fr-1 scores of the reference regions in the genome. (**A**) The TSSs of the *GAPDH* and *ATCB* genes were analyzed as active genomic regions. *Alu*, L1, and α-Satellite repeat sequences were analyzed as repressed genomic regions. The fractional proportions of the total DNA are also shown. (**B**) The ratio of the Fr-5 proportion to the Fr-1 proportion (designated as “Fr-5/Fr-1”) of the genomic regions referred in (A) was calculated and represented as a log2 value (the mean ± SD). Data obtained from at least three independent experiments were utilized for the calculations. (**C** and **D**) Percentages of the degrees of local chromatin compaction in the entire genome (C) and in TSS regions (−250 bp to ±0 bp) (D). The compaction degrees were classified into five groups based on the Fr-5/Fr-1 scores, as shown in the key.

### Genome-wide features of local chromatin compaction

To elucidate the genome-wide features of local chromatin compaction in HepG2 cells, the DNA in each fraction was analyzed by next-generation sequencing (NGS). Sequence reads were obtained as described in the MATERIALS AND METHODS. Approximately 90% of the reads from all of the fractions mapped to the human reference genome hg38 (fig. S3). All of the fractions largely consisted of intergenic and intron regions (fig. S4A and table S1), although the numbers of reads varied from 6.6 × 10^7^ (Fr-4) to 7.6 × 10^7^ (Fr-2) (fig. S4B and table S1). Hierarchical cluster analyses were performed to describe the uniqueness of Fr-1 compared with Fr-2 to Fr-5 (fig. S4C). The Fr-5/Fr-1 scores for the entire genome were calculated from the Fr-1 and Fr-5 reads. The chromatin compaction states were classified into five groups via the Fr-5/Fr-1 scores. Approximately 90% of the genomic regions were compacted at levels more than −3.5 of the Fr-5/Fr-1 scores (Fig. 3C), indicating that nearly the entire genome tends to be well compacted. However, when TSS regions were isolated, a wide variation in the Fr-5/Fr-1 scores was observed (Fig. 3D). This result suggests that the compaction level appears to be defined at the TSS of each gene.

Using a 2,000 kb region of chromosome 14, which consists of a gene-rich region at the center and relatively long intergenic regions on both sides, the local state of chromatin compaction was compared with the levels of epigenetic marks using the Integrative Genomics Viewer (Fig. 4A). The Fr-5/Fr-1 scores calculated from the Fr-1 and Fr-5 reads (Tracks 1 and 2, respectively) were represented as a heatmap (Track 3). The data for 5-methyl cytosine (5meC), DNase hypersensitivity (DHS), and histone modifications (H3K9ac, H3K27ac, H3K9me3, and H3K27me3) in HepG2 cells were obtained from the ENCODE database (accession numbers: GSM2308630, GSM2400286, GSM733638, GSM733743, GSM1003519, and GSM733754, respectively) (Tracks 4 to 9). Transcripts from the HepG2 cells used in this study were also sequenced by NGS (Track 11). As expected from Fig. 3C, there were many green stripes in Track 3 in Fig. 4A, indicating that the Fr-5/Fr-1 scores at most of the positions reached nearly −1.0. This finding suggests that the chromatin across this 2,000-kb region is compacted at a level similar to those of the repeat sequences, as shown in Fig. 3B. While 5meC was restricted in the central gene-rich region (Track 4), H3K9me3 and H3K27me3 were mainly observed outside of the central region (Tracks 8 and 9). These distributions were not correlated to the “green” regions that indicate compact chromatin (Track 3). Fig. 4B shows a magnified view of an 80 kb region from a central part of the 2,000 kb region (red bar in Track 10 of Fig. 4A). Three genes, *GEMIN2, TRAPPC6B*, and *PNN*, appear (in this order) in the 80-kb region. The *GEMIN2* and *PNN* genes are oriented rightward, while the *TRAPPC6B* gene is oriented leftward (Track 10). As marked with arrows above Track 3 in Fig. 4B, three regions with dense red stripes were found, indicating that the chromatin in these regions was poorly compacted. Intriguingly, these regions were located over the TSSs of the three genes (Tracks 3 vs. 10) and corresponded to stretches without 5meC (Tracks 3 vs. 4). These observations suggest that the absence of 5meC might be required for the local openness of chromatin at TSSs. The three regions also corresponded to stretches with DHS (Tracks 3 vs. 5), indicating that the open chromatin can be well digested by DNase I. H3K9ac, H3K27ac, and H3K27me3 were recruited over the TSSs but spread more widely compared with their distributions in the “red” regions (Tracks 3 vs. 6, 7, and 9, respectively). Again, the local chromatin compaction appeared to be distinct from any structures defined by these histone modifications.

**Fig. 4.**
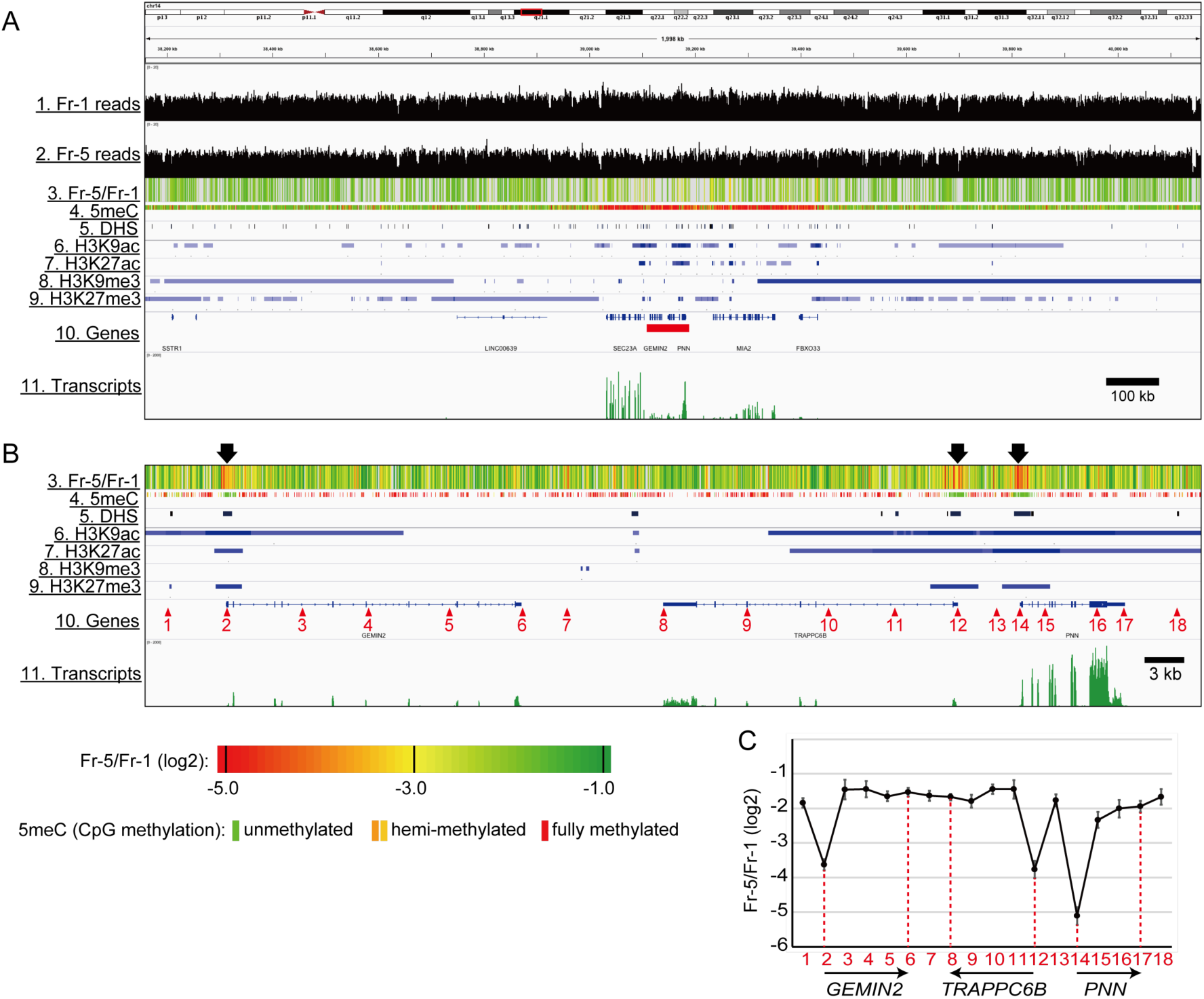
The trimmed landscape of the local chromatin compaction vs. epigenetic marks. (**A**) The Fr-5/Fr-1 magnitudes (Track 3), the epigenetic mark distributions (Tracks 4 to 9), and the transcript abundance (Track 11) in a 2,000 kb region of chromosome 14 were visualized via the Integrative Genomics Viewer. (**B**) A magnified view of an 80 kb region indicated by a red bar in Track 10 of (A). The chromatin was poorly compacted in the three regions highlighted by the arrows above Track 3. (**C**) The Fr-5/Fr-1 scores at the 18 positions indicated by the red numbered arrowheads in Track 10 of (B). Data obtained from at least three independent experiments are represented as the mean ± SD. The positions and orientations of the genes within the 80 kb region are shown with arrows.

The fractional distributions at 18 points (marked with red numbered arrowheads in Track 10 of Fig. 4B) were evaluated by qPCR (fig. S5). More than 40% of the TSSs of the three genes was found in Fr-1 (Points 2, 12, and 14), while the proportions of the other points in Fr-1 were less than 30%. The Fr-5/Fr-1 score of each point was calculated (Fig. 4C). The score at the *PNN* TSS was −5.12 (Point 14), which was lower than those of the *GEMIN2* TSS (−3.63) and the *TRAPPC6B* TSS (−3.77) (Points 2 and 12, respectively). These scores were inversely correlated to the transcript abundance, as shown in Track 11 in Fig. 4B. The regions outside the TSSs showed much higher Fr-5/Fr-1 scores (−2.33 to −1.44) (Fig. 4C), which were comparable to the scores of the repeat sequences (Fig. 3B). These results suggest that chromatin is *per se* compacted without making a distinction between repeat and non-repeat sequences; however, the compaction is locally attenuated at the TSSs of active genes.

We next focused on the bodies of active genes. To clarify the relationship between transcription level and local chromatin compaction, active genes were categorized into three groups based on their transcription levels (“Low”, “Mid”, and “High” in Fig. 5A). Nearly the entire gene bodies of the “Low” genes were evenly detected in all of the fractions, while only the TSSs were more abundant in Fr-1 and less abundant in Fr-5. This pattern was also observed for the “Mid” genes. Importantly, the Fr-1-biased distribution of the “Mid” TSSs was more apparent than that of the “Low” TSSs, suggesting an inverse correlation between the compaction at the TSS and the transcription level. On the other hand, for “High” genes, the entire bodies were largely detected in Fr-1, suggesting that openness of the chromatin across the gene body is required for a high level of transcription. Intriguingly, the transcription end sites (TESs) of the “High” genes were slightly less abundant in Fr-1 compared with the levels of the other parts of the gene bodies. Thus, moderate compaction of the TESs may be part of the transcription termination mechanism for the “High” genes. Next, the Fr-5/Fr-1 scores of the TSS and TES of each gene were calculated and plotted on scatter diagram against the transcription level. The data used for the calculations were restricted as described in the MATERIALS AND METHODS and in fig. S6A and S6B. The downward slopes of the approximation lines for both sites indicated an inverse correlation between the Fr-5/Fr-1 scores and the transcription levels (Fig. 5B). The slope of the TSS line was steeper than that of the TES line, indicating that the correlation of the TSSs was stronger than that of the TESs. We also analyzed nucleosome positioning in HepG2 cells from MNase-sequencing data in the EMBL-EBI database (accession number: E-MTAB-1750). The patterns of nucleosome occupancy around the TSS showed an NFR corresponding to a “nucleosome valley” (Fig. 5C). Consistent with previous reports (*12, 15*), a correlation between the depth of the NFR valley and the transcription level was clear when the average positioning was calculated from multiple TSSs in each group classified according to their transcription level (Fig. 5C). However, for individual genes, the depth of the valley, which was calculated as the distance between the lowest and highest nucleosome occupancy within the TSS region (−250 bp to ±0 bp), was not correlated with the transcription level (Fig. 5D). Because a relatively high transcription level occurred from TSSs with shallow NFR valley depths (less than 0.2), the absence of nucleosomes in TSSs is not necessarily a determinant for enhanced transcription. The NFR depth was also not correlated with the Fr-5/Fr-1 scores (Fig. 5E), suggesting that nucleosome eviction does not robustly influence the local chromatin compaction state.

**Fig. 5.**
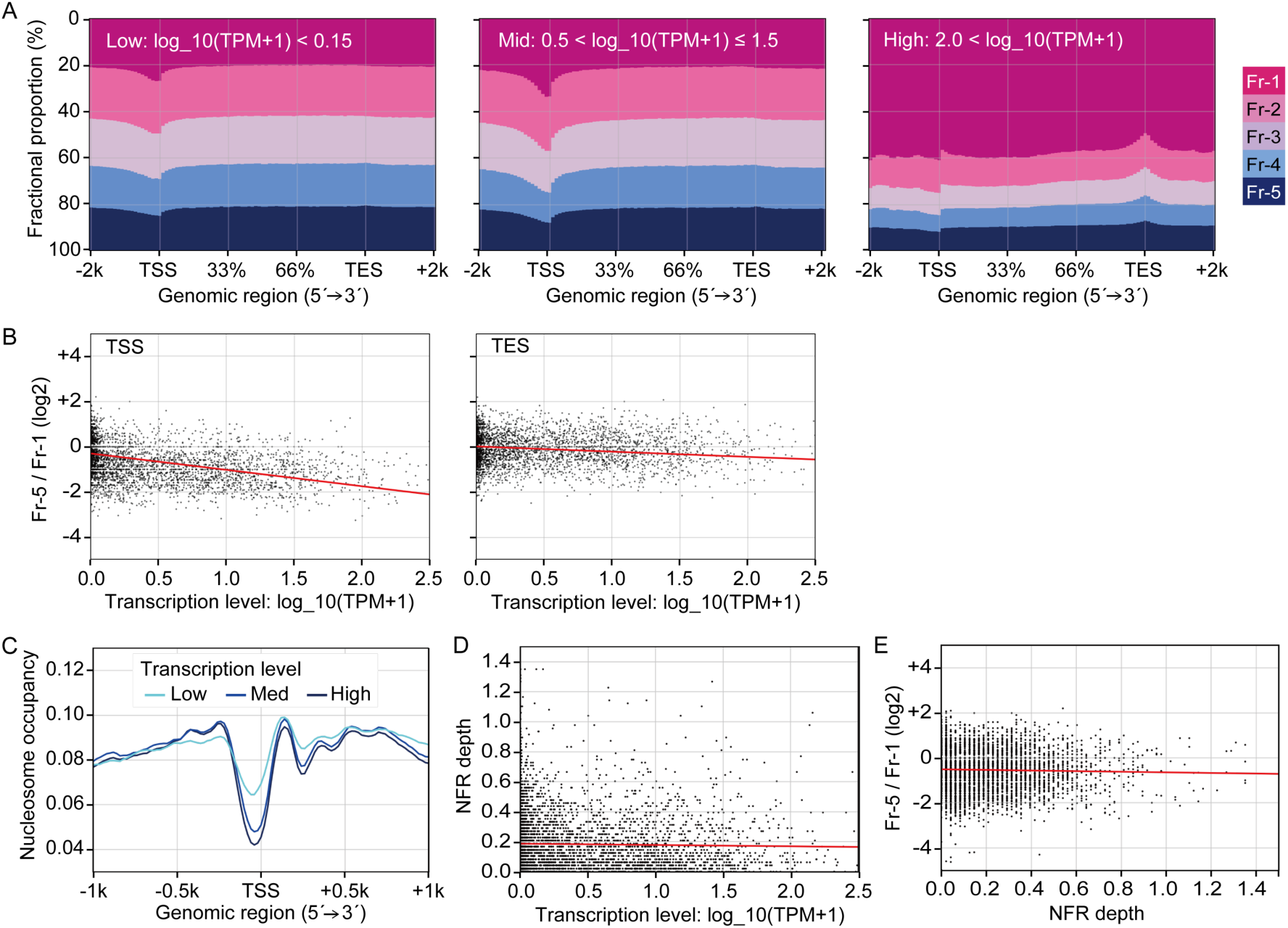
Comparison of the local chromatin compaction with the NFR formation. (**A**) The fractional proportions from 2 kb upstream of the TSSs to 2 kb downstream of the TESs of active genes. The genes were divided into three groups based on their transcription levels (values of log_10(TPM+1)). (**B**) Scatter diagrams comparing the transcription levels and the Fr-5/Fr-1 scores at the TSSs or TESs of active genes. The red line in each panel represents an approximation line of the scatter points. Data for each gene are listed in Table S2. (**C**) Nucleosome occupancy in the TSS regions (−1 kb to +1 kb) of active genes. The genes were divided to three groups as described in (A). (**D**) A scatter diagram comparing the transcription levels and NFR depth (the distance between the lowest and highest nucleosome levels within the TSS region (−250 bp to ±0 bp)). The red line represents an approximation line of the scatter points. (**E**) A scatter diagram comparing the NFR depth and the Fr-5/Fr-1 scores. The red line represents an approximation line of the scatter points.

To search for a correlation between the compaction at the TSS and RNA polymerase II (Pol II) binding, data from a chromatin immunoprecipitation-sequencing (ChIP-Seq) experiment for Pol II in HepG2 cells were obtained from the Gene Expression Omnibus (GEO) database (accession number: GSM2864932). Expectedly, the binding level of Pol II peaked at the TSSs, and the peak heights were reflected in the transcription levels (Fig. 6A). In a scatter diagram of the Pol II binding levels against the Fr-5/Fr-1 scores at the TSSs, an inverse correlation was observed (Fig. 6B), indicating that Pol II binds less frequently to TSSs in chromatin with higher Fr-5/Fr-1 scores. Taken together, these results suggest that local chromatin compaction, particularly at TSSs, can attenuate transcription possibly by reducing the binding frequency of RNA Pol II.

**Fig. 6.**
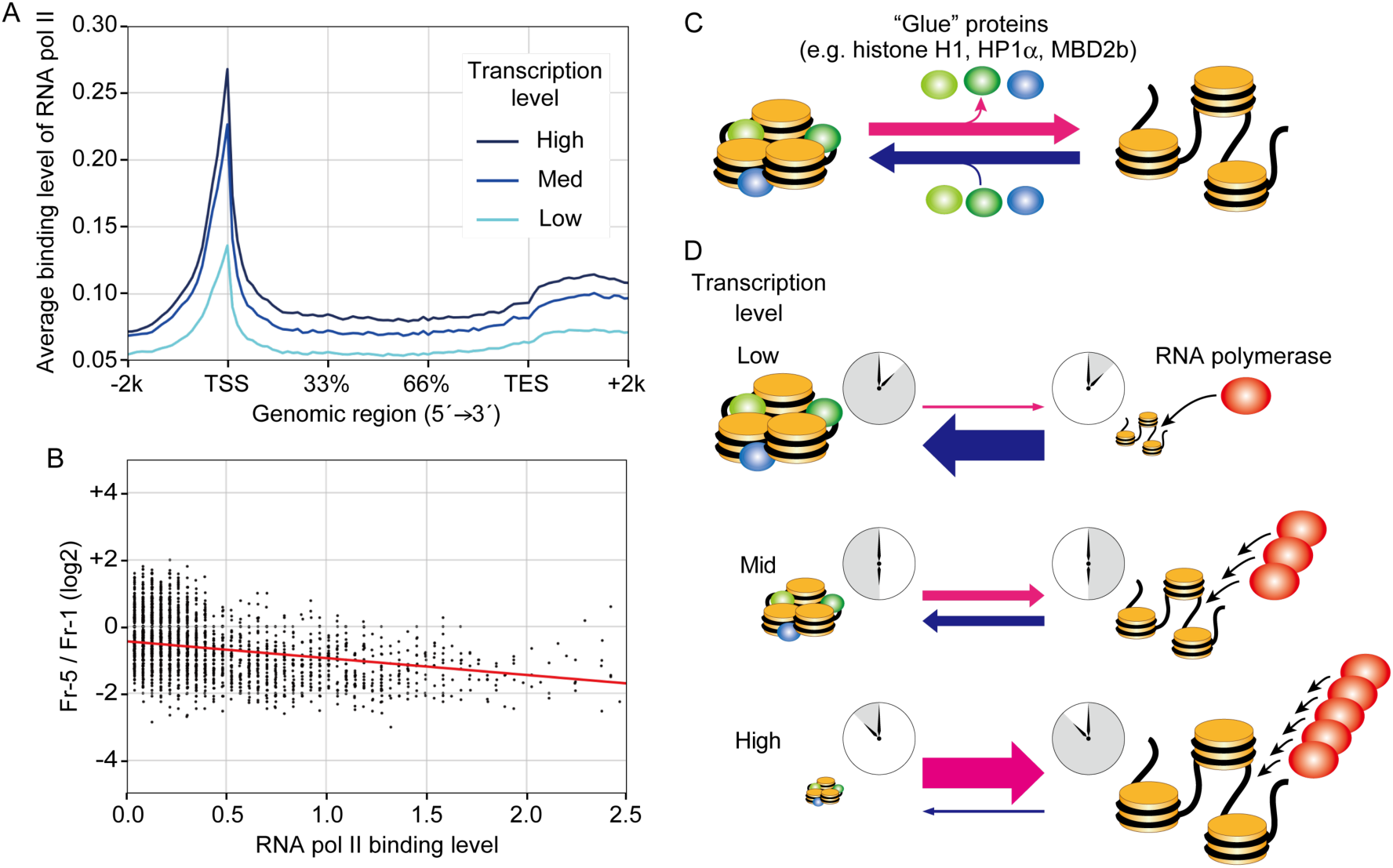
A model of the quantitative regulation of transcription by the local chromatin compaction state via the binding frequency of RNA Pol II. (**A**) The binding levels of RNA Pol II from 2 kb upstream of the TSSs to 2 kb downstream of the TESs of active genes. The genes were divided into three groups as described in Fig. 5A. (**B**) A scatter diagram comparing the RNA Pol II binding levels and the Fr-5/Fr-1 scores at the TSSs. The red line represents an approximation line of the scatter points. (**C**) Chromatin with a few nucleosomes (three in this figure) is locally compacted and opened in equilibrium. When histone H1, HP1α, and/or MBD2b are incorporated into chromatin as “glue” (green ellipses), chromatin is preferentially compacted. (**D**) The transcription level is regulated by the equilibrium in the local chromatin compaction, which affects the RNA polymerase binding frequency (red ellipses). When the equilibrium is biased toward the compact state, the time window for RNA polymerase binding is shortened (in “Low” transcription). Conversely, a bias toward the open state widens the time window for RNA polymerase binding (in “High” transcription). A middle length of the time window leads to “Mid” transcription. This model is also shown in animation videos in fig. S7.

## DISCUSSION

In the field of chromatin biology, the positioning of nucleosomes, which consist of approximately 147 bp of DNA, has been well characterized by NGS combined with MNase treatment (*36, 37*). Furthermore, an advanced 3C method, i.e., Hi-C, revealed the existence of a topologically associating domain (TAD) that includes a megabase-scale DNA element (*38–41*). Based on the data gathered using the novel approach we established in this study, we propose that local chromatin compaction represents another level in the hierarchy of chromatin structure.

DNA-processing enzymes, e.g., endonuclease or transposase, have been historically used to analyze chromatin structure (*42*). This type of analysis is based on whether the enzyme can access and then digest (or recombine with a probe sequence) the linker DNA between nucleosomes. Thus, these approaches are based on the accessibility rather than the inaccessibility of chromatin. However, in chromatin compacted within the narrow space of a few nucleosomes, the linker DNA may not be hidden among the nucleosomes, and it may be accessible to enzymes that are smaller than the RNA polymerase complex (*43*). It has been reported that endonucleases such as MNase and *Alu*I restriction enzyme can equally digest genomic DNA in open and compact chromatin, although low concentrations of such enzymes are useful for identifying hyper-accessible regions (*44, 45*). These enzyme-based methods do not seem to be sufficient for determining whether chromatin is locally compacted or not. In this study, to simultaneously evaluate the accessibility and inaccessibility of chromatin, a strategy was designed to biochemically separate open and compact chromatin via fractionation by ultracentrifugation (Fig. 1A). As confirmed by the HS-AFM experiments (Figs. 1F and 1G), chromatin was successfully fractionated according to the magnitude of its local compaction. Because this compaction is achieved based on inter-nucleosomal interactions, the absence or presence of NFRs must directly influence the formation of compact chromatin. Nevertheless, a correlation between them was not observed (Fig. 5E). Thus, local chromatin compaction appears to occur independently of the nucleosome level.

Considering fluctuating movement of nucleosomes (*26, 28–30*), chromatin is in equilibrium between moments when neighboring nucleosomes are associated with and dissociated from each other (Fig. 6C). Importantly, there is a restricted time window in this equilibrium during which RNA polymerase can access TSSs, suggesting that the length of this window could determine the RNA polymerase binding frequency. When the equilibrium is shifted toward compact chromatin, transcription would be attenuated because of less RNA polymerase binding (“Low” in Fig. 6D); when the equilibrium is shifted toward open chromatin, transcription would be enhanced because of more RNA polymerase binding (“High” in Fig. 6D) (see animation videos in fig. S7). When chromatin from a cell population was subjected to this fractionation technique, even highly active TSSs were gradationally distributed toward Fr-1 (Fig. 3), indicating that the chromatin at the TSSs is not uniformly opened. In our experiment, the equilibrium would be detected as a mixture of variously compacted structures that were obtained from a snapshot of a cell population. Because the proportion of compact vs. open structures in the mixture is reflected in the Fr-5/Fr-1 score, it would be reasonable to conclude that the Fr-5/Fr-1 score is inversely correlated to the transcription level via changes in the RNA polymerase binding frequency.

All of the epigenetic marks examined in this study, i.e., H3K9ac, H3K27ac, H3K9me3, H3K27me3, and 5meC, were detected throughout all of the fractions, although their specific fractional distributions varied slightly (Fig. 2A). These observations indicate that none of these marks are a direct determinant of whether or not chromatin is compacted. However, considering that HP1α and MBD2b were more abundant toward Fr-5 than were H3K9me3 and 5meC (Figs. 2C and 2D), some H3K9me3 and 5meC marks might have served as platforms for HP1α and MBD2b, respectively. Similarly, based on the observation of a clear enrichment of histone H1 toward Fr-5, which was not observed for histone H3 (Fig. 1E), it appears that histone H1 was recruited to a limited fraction of the nucleosomes. Thus, HP1α, MBD2b, and histone H1 may all be selectively recruited to their marks. Previous experiments with reconstituted chromatin showed that HP1α bridges two adjacent nucleosomes (*46*), and that histone H1 promotes condensation among three nucleosomes (*47*). In addition, an *in vivo* imaging study showed that histone H1 is preferentially incorporated into “clutches” of a few nucleosomes (*26*). Although the structural relationship between MBD2b and nucleosomes remains unclear because MBD2 family proteins bind directly to DNA, it is clear that HP1α and histone H1 operate on a local structure that comprises a few nucleosomes of chromatin. Histone H1 repeatedly associates with and dissociates from chromatin every few minutes (*48, 49*). Such a dynamic association of histone H1 would be related to chromatin compaction in equilibrium. Together, histone H1, HP1α, and MBD2b could function as “glue” between the adjacent nucleosomes in compact chromatin (Fig. 6C).

Heterochromatin and euchromatin, which were originally defined via cytological observations, have been proposed as intranuclear structures that act as a transcriptional switch (*50*). Furthermore, epigenetic marks have been widely recognized to influence the formation of heterochromatin and euchromatin (*51*). Our study has provided evidence of a tendency for the formation of compact versus open chromatin that extends over a few nucleosomes. These structures seem to be less temporally and spatially stable relative to heterochromatin and euchromatin. Because a portion of the H3K9me3 and 5meC marks, which are considered heterochromatin marks, were involved in local chromatin compaction via recruitment of HP1α and MBD2b, respectively, the compact chromatin might be an intermediate structure in the process leading to the formation of typical heterochromatin. Fine-tuning of transcription would be achieved at such a flexible level in the structural hierarchy of chromatin.

## MATERIALS AND METHODS

### Chromatin fractionation by using sedimentation velocity centrifugation

This chromatin fractionation technique is a modification of a method we established previously (*52, 53*), and its schema is illustrated in Fig. 1A. HepG2 cells (a human hepatoma cell line obtained from the RIKEN BRC in Japan) were used in this study. HepG2 cells were cultured in a minimum essential medium with α-modification supplemented with 10% fetal bovine serum. After washing with phosphate-buffered saline (PBS), 15 to 90 mg (wet weight) of HepG2 cells were collected into a microtube, and the concentration was adjusted to 15 mg/ml in PBS. For the crosslinking reaction, formalin (37% formaldehyde solution) was added to the cells at a final concentration of 0.5%, and the cells were agitated at room temperature for 10 min. Following addition of glycine at a final concentration of 62.5 mM to quench the formaldehyde, the cells were washed twice with ice-cold PBS, solubilized with 250 to 500 µl of Tris-based SDS lysis buffer (TSB; 1% SDS, 50 mM Tris-HCl (pH 8.0), 10 mM EDTA, and a Complete Protease Inhibitor Cocktail (Roche)), and then fragmented with a Branson Sonifier 150 (at level “2” for 5 sec 6 times). After removal of the debris using a Vivaclear Mini column (Sartorius), the cell extract, including the chromatin fragments, was layered onto an 11 ml sucrose gradient (20−60%) in chromatin dilution buffer (CDB; 1.1% Triton X-100, 0.01% SDS, 16.7 mM Tris-HCl (pH 8.0), 1.2 mM EDTA, 167 mM NaCl, and a Complete Protease Inhibitor Cocktail (Roche)) in a polyallomer centrifugation tube (Beckman Coulter). The sample was then subjected to ultracentrifugation at 256,000 x g at 4°C for 16 hrs in a Beckman SW41Ti swing rotor. Following removal of the uppermost 1.8 ml volume, five 1.8 ml fractions were collected from the top to the bottom of the tube using a micropipette.

### HS-AFM observation of fractionated chromatin

Chromatin was recovered from each fraction by immunoprecipitation with an anti-pan-histone H3 antibody (#ab1791, Abcam). Prior to the immunoprecipitation, 100 µg of the antibody was covalently conjugated to 17 mg of magnetic beads using a Dynabeads Antibody Coupling Kit (Thermo Fisher). The H3-conjugated beads (1.2 mg of beads) was mixed with each fraction, and the mixtures were agitated at 4°C overnight. After washing twice with CDB and then once with HE (50 mM Hepes (pH 7.6), 10 mM EDTA) at 4°C for 5 min each, the chromatin was eluted in 30 µl of Hepes-based SDS lysis buffer (HSB; 1% SDS, 50 mM Hepes (pH 7.6), 10 mM EDTA, and a Complete Protease Inhibitor Cocktail (Roche)). The HS-AFM observations were performed using a laboratory-built HS-AFM apparatus similar to a previously described AFM (*54*). The HS-AFM was equipped with small cantilevers (k = 0.1–0.2 N/m, f = 800– 1200kHz in solution (Olympus)) and was operated in tapping mode. The AFM styli were placed on each cantilever by electron beam deposition. A sample stage made of quartz glass was placed on the z-scanner, and a 1.5 mm diameter mica disk was glued onto the sample stage. A freshly cleaved mica surface was treated with 0.1% aminosilane for 90 s. After rinsing the surface with HE, 1.5 μl sample droplets of the chromatin preparations were placed on the mica surface and incubated for 3 min. All HS-AFM observations were performed under HE at room temperature. To estimate the sizes of the chromatin in each fraction, the diameters of the objects in the AFM images were analyzed using SPIP image analysis software (Image Metrology) and Origin (LightStone).

### Preparation of DNA from fractionated chromatin

An aliquot of each fraction corresponding to the amount of sample from 3 mg of cells was used for DNA preparation. Each aliquot was heated at 65°C overnight to reverse the crosslinking, and then successively treated with RNase A and proteinase K. Following phenol/chloroform extraction, DNA was recovered with 10 µg of glycogen by ethanol precipitation. Pellets were dissolved in 120 µl of TE (10 mM Tris (pH 7.5), 1 mM EDTA), treated with phenol/chloroform again, and then purified using a MinElute spin column (Qiagen). After elution with 30 µl of EB buffer (Qiagen), the DNA was quantified using a Quant-iT PicoGreen Kit (Thermo Fisher).

### Analyses of the proteins in the fractionated chromatin

The remaining portions of the Fr-1 to Fr-3 fractions and the Fr-4 to Fr-5 fractions were diluted with 2 volumes and 3 volumes of CDB, respectively. To recover the proteins, 100% (w/v) TCA was added to the diluted fractions at a final concentration of 20%. The mixture was chilled on ice for 30 min and then centrifuged at 21,500 × g at 4°C for 20 min. After washing with ice-cold ethanol twice, the pellets were suspended in 130 µl (Fr-0), 110 µl (Fr-1), or 50 µl (Fr-2 to Fr-5) of TCA-pellet suspension buffer (TPS; 600 mM Tris (pH 8.8), 4% SDS, 8% glycerol, 0.01% bromophenol blue). To simultaneously solubilize the pellet and reverse the crosslinking, the suspension was heated at 65°C for 24 hrs. After centrifugation at 21,500 × g at 4°C for 10 min, the proteins were recovered in the supernatants. To observe the total protein in each fraction, the volumes of the protein preparations were adjusted with TPS among the fractions. Following treatment with 100 mM dithiothreitol (DTT) at 100°C for 5 min, the total protein was size-separated on an 8% polyacrylamide gel and stained with SYPRO Ruby (Lonza). When the contents of the protein preparation were analyzed by western blotting, the preparations were adjusted among the fractions with TPS based on the amount of DNA. Following treatment with DTT, the protein preparations were loaded onto a 10% (for histones) or 8% (for non-histone proteins) SDS-PAGE gel, ran, and then transferred to a nitrocellulose membrane (0.2 µm pore size). After blocking in 5% skim milk in Tris-buffered saline (TBS) with 0.1% Tween 20, the membranes were sequentially exposed to a primary antibody, a biotinylated secondary antibody, and streptavidin-conjugated alkaline phosphatase (GE Healthcare). They were then developed with a BCIP-NBT Solution Kit (Nacalai). To estimate the fractional distributions of the proteins, a standard curve for quantitation was calculated from the blot signals from serially diluted samples, whose intensities were measured using ImageJ. The primary and secondary antibodies are listed in Table S3.

### Analyses of 5meC in the DNA from the fractionated chromatin

Two hundred ng of the DNA (adjusted to 30 µl) from each fraction was denatured by heating at 100°C for 5 min. After being immediately chilled on ice for 5 min, the DNA was spotted onto a nitrocellulose membrane (0.2 µm pore size) using a Bio-Dot Apparatus (Bio-Rad). After the membrane was baked at 80°C for 2 hrs, immunoblotting with an anti-5-methyl cytosine antibody (#ab1884, Abcam) was performed as described in the previous section.

### Analyses of the DNA from the fractionated chromatin

For qPCR analyses, 500 pg (for the non-repeat sequences), 62.5 pg (for the L1 sequence), or 16.7 pg (for the *Alu* and α-satellite sequences) of the recovered DNA was used for a single reaction. To generate a standard curve, serially diluted human genomic DNA (0.76−12,500 pg; #D4642, Sigma-Aldrich) was utilized as previously described (*53, 55*). The amount of each sequence was estimated from the respective PCR cycle threshold (Ct) value plotted on the standard curve. A 1:3 mixture of a QuantiFast SYBR Green PCR Kit (Qiagen) and a FastStart SYBR Green Master (Roche) in a real-time PCR machine (#7900HT, Applied Biosystems) was used for the qPCR. The PCR primers are listed in Table S4. For preparation of an NGS sequence library of the DNA from the fractionated chromatin, 28 ng of the DNA in a Crimp-cap microTUBE (Covaris) was fragmented with an LE220 Focused-ultrasonicator (Covaris). The configuration of the ultrasonication process was as follows: temperature, 7°C; duty factor, 30%; peak incident power, 450 W; cycles per burst, 200; and time, 190 sec. Following concentration via a DNA Clean & Concentrator-5 (Zymo Research), the fragmented DNA was converted to a sequence library using a KAPA Hyper Library Preparation Kit (KAPA Biosystems). To analyze transcripts in the HepG2 cells, 2 µg of total RNA was converted to a sequence library using a KAPA Stranded mRNA-seq Kit (KAPA Biosystems). These libraries were analyzed using a HiSeq 2500 sequencer (Illumina) with the following specifications: Read1, 50 cycles.

### Bioinformatic analyses

The sequence reads were trimmed via the fastx_trimmer function of the FASTX-toolkit (version: 0.0.14), retaining the last 50 bps (parameter: “-l 50”). HISAT2 (version 2.0.4) was used to map the reads to the human hg38 genome with the default parameters. Samtools-0.1.19 fulfilled the requirement of HISAT2. The reads employed for our analyses were qualified via Samtools (version 1.3) with “samtools view -q 4” and were confirmed to not overlap in repetitive sequences of hg38 via intersectBed (bedtools v2.25.0) with option -v. The repetitive sequence data were obtained from the UCSC Genome Browser (https://genome.ucsc.edu/cgi-bin/hgTables). The transcription levels were evaluated as transcripts per kilobase million (TPM) using Sailfish (beta v0.10.0). Using this parameter, the transcription levels were categorized as follows: “Low”, log_10(TPM+1) < 0.15; “Mid”, 0.5 < log_10(TPM+1) <= 1.5; “High”, 2.0 < log_10(TPM+1). Ensemble76 reference data with the option “-p 20 -l SR -r” were employed for the transcript annotation. Python script “read_distrobution.py” in RSeQC (version 2.6.4) was used to count the reads mapped to intergenic or intragenic regions, as shown in figs. S4A and S4B. Hg38 genome annotation data (hg38_Gencode_V23.bed file in Sourceforge (https://sourceforge.net/p/rseqc/activity)) was used with the -r option of “read_distrobution.py”. Hierarchical clustering of the composition ratios of each fraction was performed using the pvclust package (http://stat.sys.i.kyoto-u.ac.jp/prog/pvclust/) of the R program (with the bootstrap trial time equal to 1,000 (nboot = 1,000)). The read depth analyses were performed using “bam2wig.py” in RSeQC (version 2.6.4), specifying the wigsum as 8500000000 (-t 8500000000), skipping non-unique hits reads (-u), and fixing the chromosome sizes (-s “hg38 chromosome size file”). To obtain the Fr-5/Fr-1 scores for the entire genome, WiggleTools (https://github.com/Ensembl/WiggleTools) was used. First, the read depth scores were scaled by the amount of recovered DNA (“wiggletools scale” with Table S5), and 0.001 was added to each of the scores (“wiggletools offset 0.001”, to avoid substitution 0 for logarithm operation (Fr-5/Fr-1)). Next, division and logarithm operations (to obtain the Fr-5/Fr-1 scores) were performed (“wiggletools ratio” and “wiggletools log 2”). A list of genes and gene predictions was downloaded from the UCSC Genome Browser (https://genome.ucsc.edu/cgi-bin/hgTables) to obtain the coordinates of the TSS regions (−250 bp to ±0 bp) for Fig. 3D. To obtain the fractional proportions of Fr-1 to Fr-5 at the TSSs and TESs, the R script “ngs.plot.r” of the ngs.plot package (https://github.com/shenlab-sinai/ngsplot) was employed. The data in Fig. 5A were obtained from the results of “ngs.plot.r” with the option “-G hg38 -R ‘genebody’”. The genes that satisfied the requirement that the “read count per million mapped” values at the TSSs and TESs were larger than 0.05 (to avoid substitution 0 for logarithm operation of the Fr-5/Fr-1) were used to extract the qualified data (see figs. S6A and S6B), as listed in Table S2, and to generate the scatter plots, along with the calculated approximation lines, presented in Fig. 5B. MNase-sequencing data were downloaded from the EMBL-EBI database (accession number : E-MTAB-1750, ERR325293). The format of the data was converted as csfastq to fastq using the Perl script “csfq2fq.pl” (obtained from https://gist.github.com/pcantalupo). Bowtie2 (version 2-2.3.5.1) was used to map the reads to the human hg38 genome with the default settings. A ChIP-Seq dataset (annotated to hg19) for RNA Pol II in HepG2 cells were downloaded from the GEO database. The data were re-annotated to hg38 using the liftOver (https://genome.ucsc.edu/cgi-bin/hgLiftOver). The protocols for Figs. 5C and 6A, and Figs. 5D, 5E, and 6B were the same as the analyses for Figs. 5A and 5B, respectively.

## SUPPLEMENTARY MATERIALS

Fig. S1. The size distributions of the DNA fragments prepared from the fractionated chromatin using various formaldehyde concentrations.

Fig. S2. The fractional proportions of the TSSs of the *GAPDH* (active) gene and the *IL2RA* (repressed) gene in the fractionated chromatin following treatment with various formaldehyde concentrations.

Fig. S3. The percentages of the mapped vs. unmapped reads in each fraction. Fig. S4. Annotation analyses of the NGS reads.

Fig. S5. The fractional proportions of the 18 positions marked by red numbered arrowheads in Fig. 4B.

Fig. S6. Establishing a cut-off point for the Fr-5 and Fr-1 reads (per million mapped reads) for the scatter diagrams in Fig. 5.

Fig. S7. Animation videos showing dynamics of local chromatin compaction at TSSs for controlling transcription levels.

Table S1. NGS read count of the DNA obtained from each fraction.

Table S2. List of genes annotated in Fig. 5B.

Table S3. List of antibodies used in this study.

Table S4. List of PCR primers used in this study.

Table S5. Data list of the NGS reads in each fraction.

## Acknowledgments

We thank Akihiro Matsushima and Manabu Ishii for their assistance with the infrastructure for the data analyses in the Laboratory for Bioinformatics Research, RIKEN Center for Biosystems Dynamics Research. The antibodies against H3K9un and H3K27un were gifts from the MAB Institute, Inc.

## Author contributions

S.I. conceptually designed this study and wrote the manuscript. S.I., N.K., and Y.Sh. performed and analyzed the biochemical experiments, including the chromatin fractionation. Y.Sa., M.U., and I.N. performed the NGS experiments. T.K. and I.N. designed and performed the bioinformatic analyses. H.Y. and M.A. performed and analyzed the HS-AFM experiments.

